# Polycomb regulation is coupled to cell cycle transition in pluripotent stem cells

**DOI:** 10.1101/2020.01.15.907519

**Authors:** Helena G. Asenjo, Amador Gallardo, Lourdes López-Onieva, Irene Tejada, Jordi Martorell-Marugán, Pedro Carmona-Sáez, David Landeira

**Author notes:** Correspondence to David Landeira. **Non-specialist teaser** Epigenetic regulation by the chromatin modifier Polycomb is modulated across cell cycle in stem cells.

## Abstract

When self-renewing pluripotent cells receive a differentiation signal, ongoing cell duplication needs to be coordinated with entry into a differentiation program. Accordingly, transcriptional activation of lineage specifiers genes and cell differentiation is confined to the G1-phase of the cell cycle by unknown mechanisms. We found that Polycomb repressive complex 2 (PRC2) subunits are differentially recruited to lineage specifier gene promoters across cell cycle in mouse embryonic stem cells (mESCs). Jarid2 and the catalytic subunit Ezh2 are dramatically accumulated at target promoters during S and G2, while the transcriptionally activating subunits EPOP and EloB are enriched during G1. Importantly, fluctuations in the recruitment of PRC2 subunits promote changes in RNA synthesis and RNA polymerase II binding that are compromised in Jarid2 -/- mESCs. Overall, we show that differential recruitment of PRC2 subunits across cell cycle enables the establishment of a chromatin state that facilitates the induction of cell differentiation in G1.

## Introduction

Deciphering the molecular mechanisms regulating pluripotent stem cell differentiation is of fundamental importance to understand mammalian development and for safe application of pluripotent stem cell-based therapies (*1*). Self-renewing mouse embryonic stem cells (mESCs) can be derived from the developing mouse blastocyst and provide a well-established system to study the molecular basis of pluripotency and early development. mESCs can differentiate into all cell types of the adult organism. However, the ability of individual mESCs within the population to respond to differentiation stimuli can be markedly different (*2-4*). This reveals a key aspect of the regulation of pluripotent cell differentiation and poses an important handicap for the application of stem-cell-based therapies to humans. The features determining the responsiveness of individual cells to differentiation cues are mostly unknown and are currently subject of intense debate. Accumulated evidence indicates that transcriptional activation of lineage specifiers genes and cell differentiation in pluripotent cells is confined to the G1-phase of the cell cycle (*5-8*). This observation partly explains the observed functional heterogeneity of pluripotent cell populations and highlights a very important regulatory feature of stem cell differentiation. Despite its obvious relevance, very little is known about the molecular mechanisms underlying this type of regulation (*9*).

Polycomb group (PcG) proteins are a hallmark of epigenetic control in eukaryotes and key regulators of mammalian development, cancer progression and stem cell differentiation (*10, 11*). In mESCs, PcG proteins associate to form Polycomb Repressive Complexes 1 and 2 (PRC1 and PRC2) that catalyse H2AK119 monoubiquitination and H3K27 methylation respectively (*11*). PRCs bind and repress hundreds of developmental regulator genes that will be activated later during cell differentiation (*12, 13*). In mESCs, PRC-target repressed genes display nucleosomes modified with functionally opposing histone modifications including H3K4me3 and H3K27me3 (hence the designation of bivalent chromatin), leaky production of transcripts and binding of poised RNAPII phosphorylated on Ser5 (Ser5-RNAPII) (*14-17*). PRC2 is typically composed by core subunits Eed, Suz12, Rbbp4/7 and Ezh1/2 of which the last one harbours the histone methyltransferase catalytic function (*11*). PRC2 function is regulated by non-stoichiometric accessory subunits that are differentially expressed during development (*11*). mESCs express high levels of Jarid2 that recruits and enhances PRC2 activity (*11, 18-22*). Additionally, mESCs express the PRC2-interacting protein EPOP, that mediates recruitment of EloB and promote low-level transcription of bivalent genes (*23*). It’s been proposed that Jarid2 and EPOP form mutually exclusive complexes (*23*). In agreement, studies of PcG proteins interactome have shown the existence of at least two types of PRC2 subcomplexes, one containing EPOP and PCL1-3, and another one containing Jarid2 and Aebp2 (*11*). They have been proposed to be termed PRC2.1 and PRC2.2 respectively (*24*). Importantly, the functional relevance of PRC2 subcomplexes specialization remains unknown.

In this study, we asked whether chromatin regulation by PRC2 was linked to the preference of pluripotent cells to enter differentiation in G1. We found that recruitment of PRC2.1 complex to target promoters is favoured in G1 while binding of PRC2.2 complex increases during S and G2-M leading to gradual accumulation of the catalytic subunit Ezh2 at bivalent promoters. This is accompanied by enhanced gene repression and accumulation of paused Ser5-RNAPII at bivalent promoters during S, G2-M. This cell cycle-dependent regulation is particularly evident at pioneering lineage specifiers, whose tight regulation is hindered in Jarid2 -/- mESCs. Taken together, our results strongly suggest that differential recruitment of PRC2 complexes across cell cycle is key to establish a chromatin state that facilitates the induction of cell differentiation in G1.

## Results

### Recruitment of Ezh2 to bivalent promoters increases during S, G2-M phases

We established wild-type mESCs that stably express the Fluorescent Ubiquitination Cell Cycle Indicator (Fucci) reporter system (*25*) (Fucci-mESCs) (Fig. S1A), and used flow cytometry to isolate highly enriched populations of cells in G1-phase (83+/-2%), S-phase (55+/-1%) and G2-M-phases (81+/-1%) (Fig. 1A). Genome wide analysis of Ezh2 binding by chromatin immunoprecipitation followed by sequencing (ChIP-seq) readily revealed the prevalence of recruitment of Ezh2 to chromatin in S, G2-M compared to G1 (Ezh2 binding peaks: 6304 in G1, 9615 in S and 15950 in G2-M). Using published data, we established a list of high confidence Polycomb-target bivalent genes (HC-bivalent, n=1678) (Fig. 1B). As controls, we used transcriptionally active (n=1557) and hypermethylated (n=656) genes that are not targeted by PRC2 (see methods). Heatmap analysis of Ezh2 binding to HC-bivalent genes showed that recruitment of Ezh2 was increased as cells exit G1 and transit into S and G2-M phases (Fig. 1C). Comparison of Ezh2-binding at HC-bivalent gene promoters showed that although Ezh2 accumulates around the Transcription Start Site (TSS) of bivalent genes at all cell cycle phases, the amount of Ezh2 bound gradually increases as cells exit G1 and transit through the cell cycle (Fig. 1D,E, S1B,C). Recruitment of Ezh2 in G1 appeared weak compared to G2-M (Fig. 1D,E), but it was evident when compared to hypermethylated promoters known to be devoid of PRC2 (Fig. 1F, Fig. S1D). Importantly, analysis of Ezh2-binding at individual promoters revealed a very consistent and gradual accumulation of Ezh2 during S and G2-M in most (1576 out of 1677, 93.9%) HC-bivalent gene promoters (see cluster I and cluster II in Fig. 1G, Fig. S1E) including the archetypical *Hoxd* gene cluster (Fig. 1H). These observations were confirmed by ChIP-qPCR for Ezh2 and analysis of a subset of well-characterized (*18*) PRC2-target promoters (Fig. 1I). Thus, we concluded that binding of Ezh2 to target promoters is dramatically enhanced upon G1 exit in mESCs.

**Figure 1.**
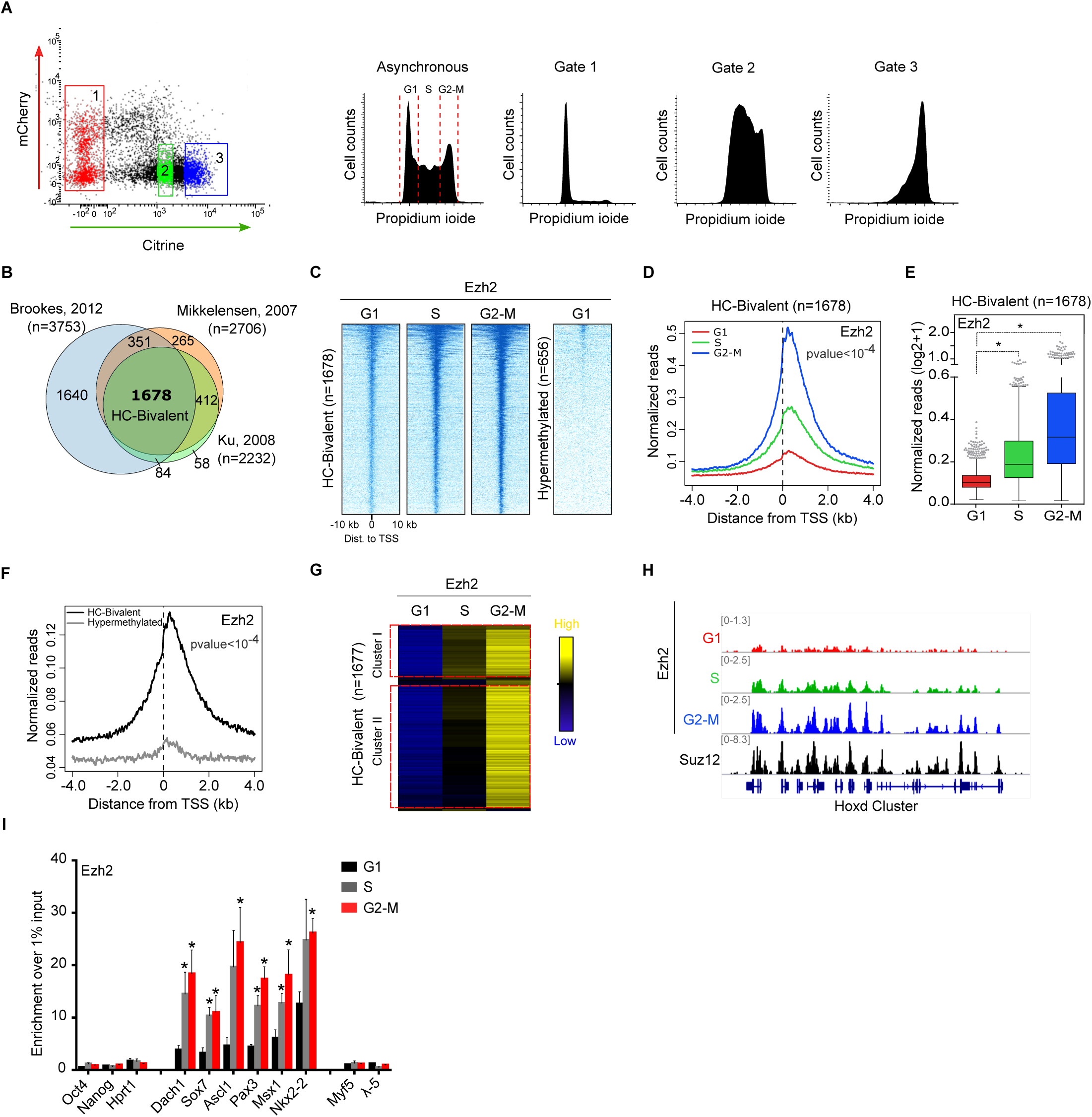
Recruitment of the PRC2-core subunit Ezh2 to bivalent genes increases during S, G2-M phases of the cell cycle. **(C)** Heatmaps of normalized Ezh2 ChIP-seq reads around the TSS of HC-bivalent promoters at different cell cycle phases. Heatmap of hypermethylated promoters is shown as a negative control. **(D)** Average binding profile of Ezh2 around the TSS of HC-bivalent promoters in G1 (red), S (green) and G2-M (blue). **(E)** Quantification of Ezh2-binding signal at the promoter regions (−0.5 kb to +1.5 kb relative to TSS) of HC-bivalent genes in indicated cell cycle phases. **(F)** Average binding profile of Ezh2 around the TSS of HC-bivalent (black) and hypermethylated (grey) promoters in G1. **(G)** Hierarchical clustering analysis of binding of Ezh2 to the promoter region (−0.5 kb to +1.5 kb relative to TSS) of HC-bivalent genes at indicated phases of the cell cycle. Binding relative to the average is presented. **(H)** Genome browser view of Ezh2 ChIP-seq data across cell cycle at the *Hoxd* gene cluster. Suz12 binding was analysed using published data (*19*). **(I)** Histogram showing enrichment of Ezh2 to PRC2-target promoter regions (*Dach1, Sox7, Ascl1, Pax3, Msx1, Nkx2-2*) in G1 (black), S (grey) and G2-M (red) assayed by ChIP-qPCR. Active (*Oct4, Nanog, Hprt1*) and hypermethylated (*Myf5, λ-5*) gene promoters were used as negative controls. Mean ± SEM of three experiments is shown. **E,I**. * marks statistically significant differences.

### Inverse binding patterns of Jarid2 and EPOP across cell cycle

We next asked whether accumulation of Ezh2 at bivalent promoters during S-G2-M reflected increased binding of PRC2.1, PRC2.2 or both. ChIP-seq analysis of Jarid2 binding in cell cycle-sorted Fucci-mESCs demonstrated that recruitment of Jarid2 to the promoter region of HC-bivalent genes was augmented in S and G2-M compared to G1 (Fig. 2A, B, Fig. S2A, B). Increased recruitment of Jarid2 to target genes in S, G2-M was evident but quantitatively less accused than changes found in binding of Ezh2 across cell cycle (Fig. 1D). Notwithstanding, clustering analysis showed a very consistent tendency to accumulate Jarid2 in S-G2-M at individual HC-bivalent genes (1404 out of 1677, 83.7%) (see cluster I and cluster II in Fig. 2C). Jarid2 was bound to most of Ezh2-bound HC-promoters in G2-M (1188 out of 1262, 94%) (Fig. S2C) and binding of Jarid2 and Ezh2 around the TSS of HC-bivalent gene promoters in G2-M displayed concordant distributions (Fig. 2D). Importantly, correlation analysis demonstrated that accumulation of Jarid2 at target genes in S, G2-M correlates with increased recruitment of Ezh2 to the same promoter regions (Fig. 2E). Coordinated recruitment of Jarid2 and Ezh2 to target genes upon G1 exit was also observed at individual candidate regions (i.e. gene *Adra2c*) (Fig. 2F).

**Figure 2.**
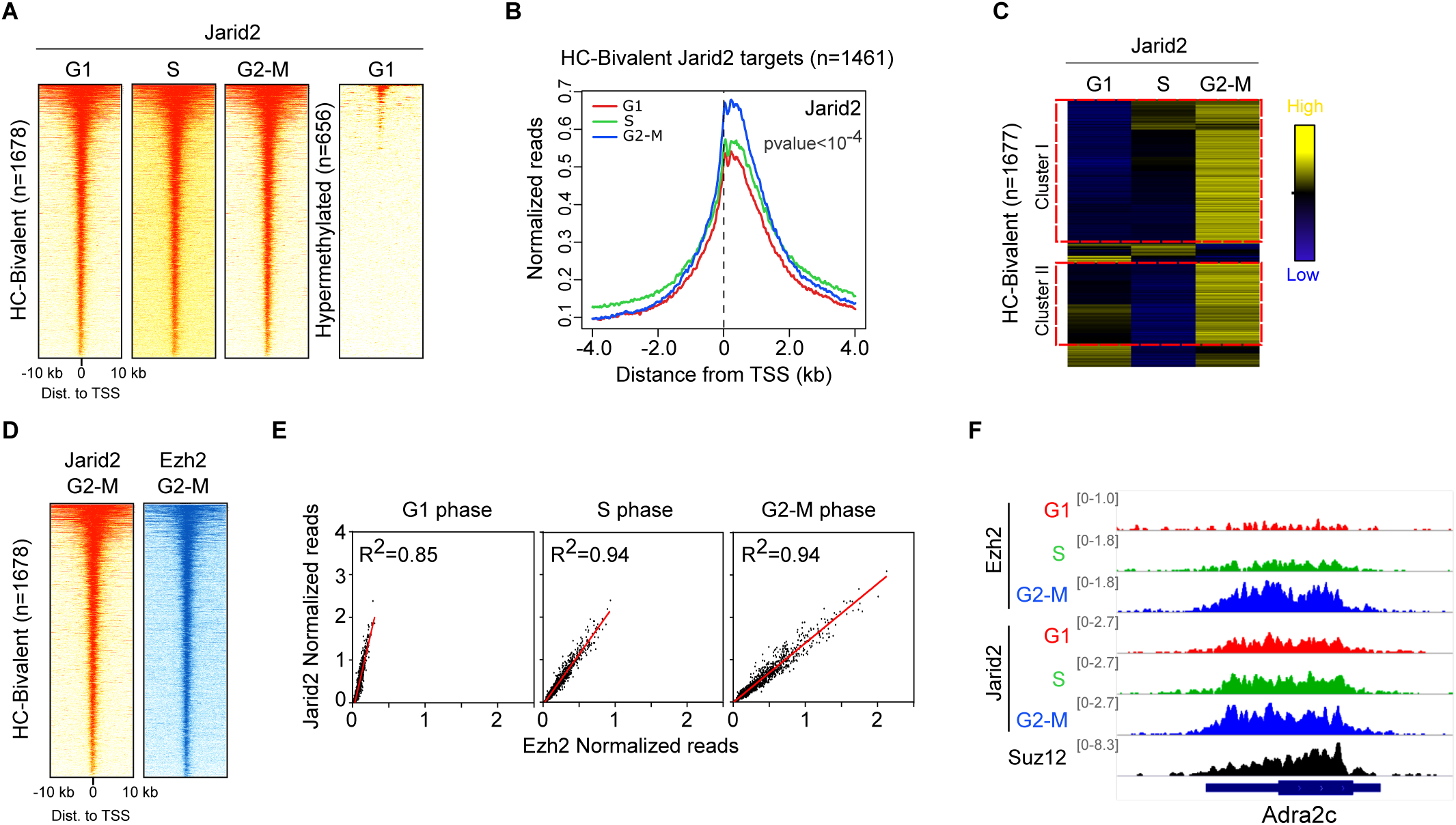
Binding of Jarid2 to target promoters increases during S, G2-M phases of the cell cycle. **(A)** Flow cytometry dot plot analysis of Fucci-mESCs indicating sorting gates used to obtain cell populations enriched in G1 (Gate1), S (Gate 2) and G2-M (Gate 3) cell cycle phases (left panel). Sorted cells were stained with propidium ioide and analysed by flow cytometry (right panel). **(B)** Venn diagrams of bivalent genes previously published in (*36-38*). **(A)** Heatmaps of normalized ChIP-seq reads showing the binding of Jarid2 around the TSS of HC-bivalent promoters at different cell cycle phases. Heatmap of hypermethylated promoters is shown as a negative control. **(B)** Average binding profile of Jarid2 around the TSS of HC-bivalent promoters in G1 (red), S (green) and G2-M (blue). **(C)** Hierarchical clustering analysis of binding of Jarid2 to the promoter region (−0.5 kb to +1.5 kb relative to TSS) of HC-bivalent genes at indicated phases of the cell cycle. Binding relative to the average is presented. **(D)** Heatmaps comparing the binding of Jarid2 and Ezh2 around the TSS of HC-bivalent promoters in G2-M. **(E)** Linear regression analysis showing the correlation between the binding signals of Jarid2 and Ezh2 at HC-bivalent promoters (−0.5 kb to +1.5 kb relative to TSS) at indicated cell cycle phases. **(F)** Genome browser view of Ezh2 and Jarid2 binding across cell cycle at the *Adra2c* bivalent gene. Suz12 binding was analysed using published data (*19*).

We next analysed binding of the PRC2.1-subunit EPOP. In stark contrast to the binding pattern observed for Ezh2 and Jarid2, binding of EPOP around the TSS of HC-bivalent genes was reduced in cells in G2-M (Fig. 3A-C, Fig. S3A, B). Clustering analysis showed that increased binding of EPOP in G1 compared to G2-M can be observed in most (1426 out of 1677, 85%) bivalent gene promoters (see cluster I in Fig. S3C). Increased recruitment of EPOP to target promoters in G1 was also observed by ChIP-qPCR for a representative set of PRC2-target promoters (Fig. 3D). We next addressed whether increased binding of EPOP in G1 led to augmented recruitment of EloB in this cell cycle phase. As expected, ChIP-seq analysis showed that EloB is recruited to HC-bivalent genes more profoundly in G1 compared to G2-M in average (Fig. 3E-G, Fig. S3D,E) and individual gene (Fig. S3F) analyses. Importantly, global levels of Ezh2, Jarid2 and EPOP across cell cycle were unchanged (Fig. 3H) indicating that differential recruitment to target genes is regulated by changes in protein interactions rather than changes in protein abundance. Taken together, these results demonstrate that Jarid2 and Ezh2 recruitment is increased during S, G2-M while EPOP and EloB are preferentially bound to chromatin in G1. This indicates that PRC2.1 complex is preferentially recruited to target promoters during G1 while PRC2.2 complex is favoured during S, G2-M phases.

**Figure 3.**
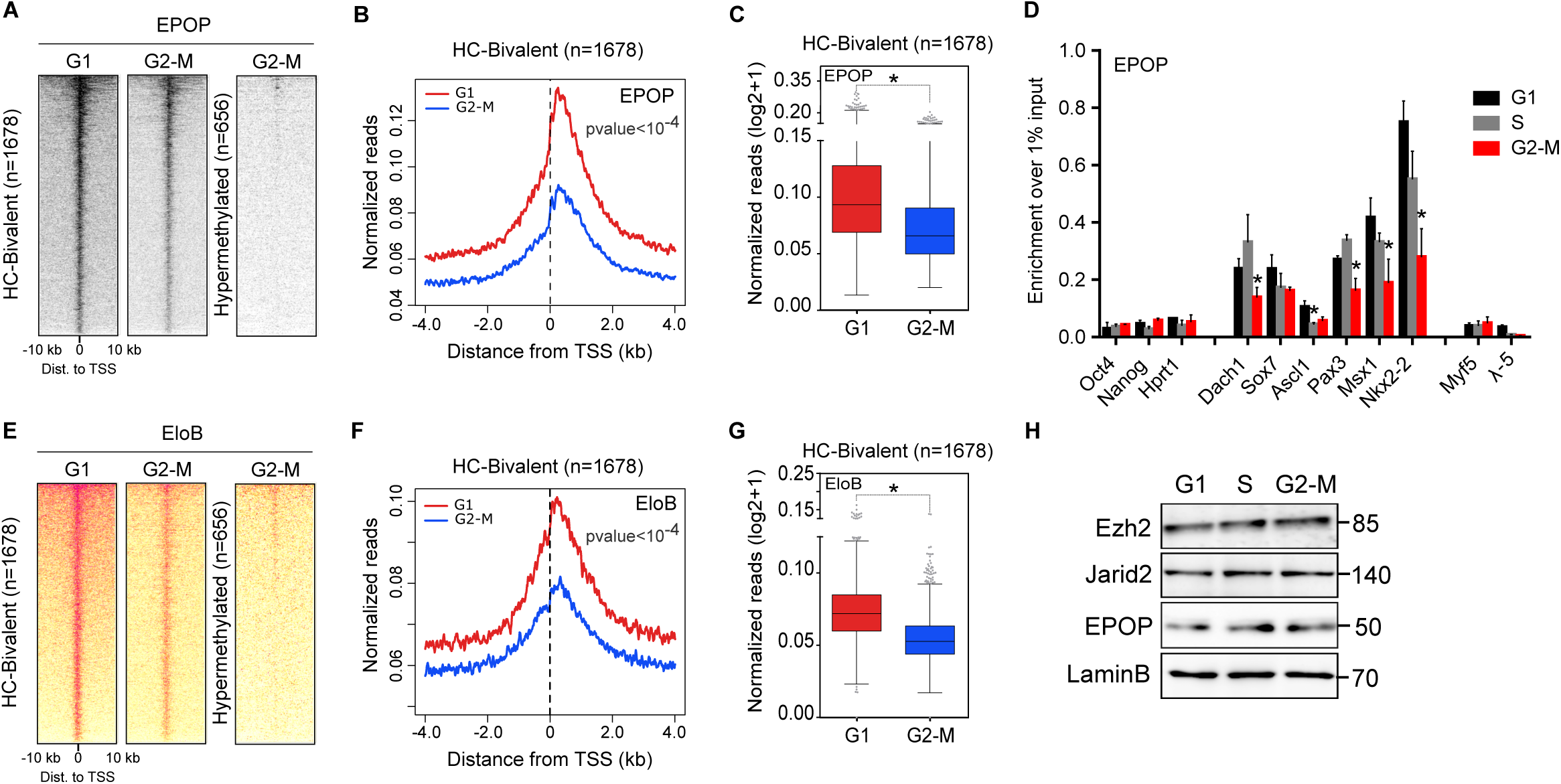
Binding of EPOP and EloB to bivalent promoters is enhanced in G1. **(A)** Heatmaps showing the binding of EPOP around the TSS of HC-bivalent promoters in G1 and G2-M. Heatmap of hypermethylated promoters is shown as a negative control. **(B)** Average binding profile of EPOP around the TSS of HC-bivalent promoters in G1 (red) and G2-M (blue). **(C)** Boxplot of EPOP-binding signal at the promoter regions (−0.5 kb to +1.5 kb relative to TSS) of HC-bivalent genes in indicated cell cycle phases. **(D)** Histogram showing enrichment of EPOP to PRC2-target promoter regions (*Dach1, Sox7, Ascl1, Pax3, Msx1, Nkx2-2*) in G1 (black), S (grey) and G2-M (red) assayed by ChIP-qPCR. Active (*Oct4, Nanog, Hprt1*) and hypermethylated (*Myf5, λ-5*) gene promoters were used as negative controls. Mean ± SEM of three experiments are shown. **(E)** Heatmaps showing the binding of EloB around the TSS of HC-bivalent promoters in G1 and G2-M. Heatmap of hypermethylated promoters is shown as a negative control. **(F)** Average binding profile of EloB around the TSS of HC-bivalent promoters in G1 (red) and G2-M (blue). **(G)** Quantification of EloB-binding signal at the promoter regions (−0.5 kb to +1.5 kb relative to TSS) of HC-bivalent genes in indicated cell cycle phases. **(H)** Whole cell lysate western blots comparing Ezh2, Jarid2 and EPOP protein levels in G1, S and G2-M. Lamin B was used as a loading control. **C, D, G**. * marks statistically significant differences.

### RNA synthesis is reduced and Ser5-RNAPII is accumulated at PRC2-target genes during S, G2-M

We next questioned whether changes in recruitment of PRC2 subunits across cell cycle had functional consequences in the transcriptional regulation of target genes. To analyse low-level RNA transcription typically found at bivalent promoters, we used 4-thiouridine (4sU)-tagging followed by high throughput sequencing (4sU-seq). 4sU-seq permits the analysis of newly transcribed RNA (*26, 27*) (Fig. S4A) and thus allowed us to minimize contamination of RNA molecules produced in previous cell cycle phases. We found increased production of RNA in G1 compared to S-G2-M at HC-bivalent genes but not at promoters of active genes (Fig. 4A, B and Fig. S4B, C). Out of 1655 HC-bivalent genes, 974 genes showed differences in RNA production between G1 and G2-M. Among these, most of them (715, 73.4%) were downregulated in G2-M compared to G1 (Fig. S4D) revealing a consistent tendency of bivalent genes to be more strictly repressed during S, G2-M than in G1. In agreement, analysis of individual candidate genes confirmed that bivalent promoters (i.e. *Nes*) are overtly repressed in S, G2-M (Fig. 4C). Importantly, level of RNA synthesised from bivalent promoters in G1 was still very low compared to the level of RNA produced at active promoters (compare scale of y-axis in HC-Bivalent and Active plots, Fig. 4A), indicating that transient alleviation of PRC2-repression in G1 results in increased leaky transcription rather than full activation of bivalent genes.

**Figure 4.**
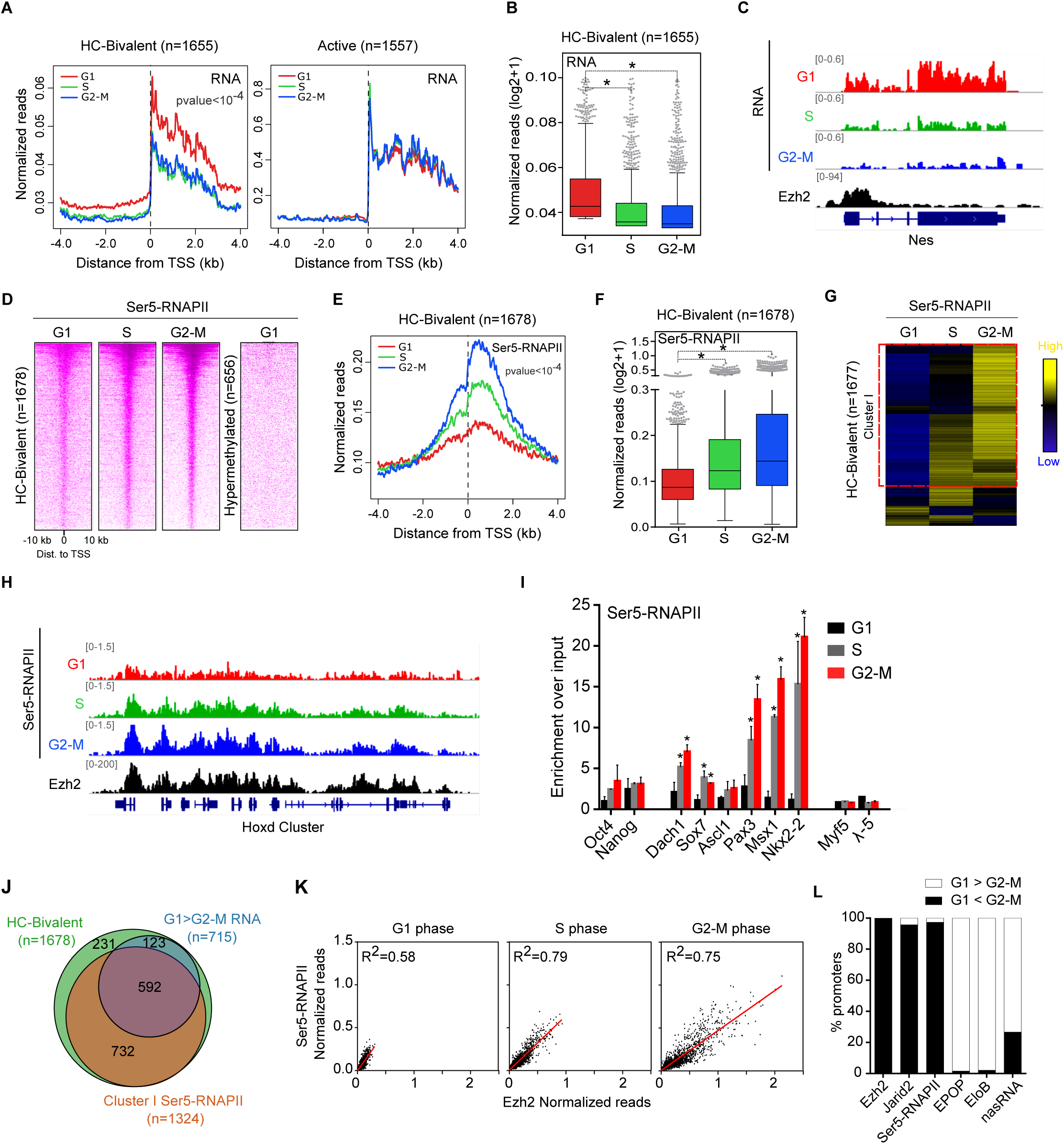
RNA synthesis is downregulated and Ser5-RNAPII is accumulated at PRC2-target promoters during S, G2-M. **(A)** Average RNA production from HC-bivalent (left panel) and active (right panel) promoters in G1 (red), S (green) and G2-M (blue). **(B)** Boxplot comparing 4sU-seq reads mapped to the proximal promoter region (TSS to +3Kb) of HC-bivalent genes in indicated cell cycle phases. **(C)** Genome browser view of RNA synthesis at indicated cell cycle phases at the bivalent gene *Nes*. Ezh2 binding was analysed using published data (*46*). **(D)** Heatmaps showing the binding of Ser5-RNAPII around the TSS of HC-bivalent promoters in G1, S and G2-M. Heatmap of hypermethylated promoters is shown as a negative control. **(E)** Average binding profile of Ser5-RNAPII around the TSS of HC-bivalent gene promoters in G1 (red), S (green) and G2-M (blue). **(F)** Quantification of Ser5-RNAPII-binding signal at the promoter regions (−0.5 kb to +1.5 kb relative to TSS) of HC-bivalent genes in indicated cell cycle phases. **(G)** Hierarchical clustering analysis of binding of Ser5-RNAPII to the promoter region (−0.5 kb to +1.5 kb relative to TSS) of HC-bivalent genes at indicated phases of the cell cycle. Binding relative to the average is presented. **(H)** Genome browser view of the binding of Ser5-RNAPII across cell cycle at the *Hoxd* gene cluster. Ezh2 binding was analysed using published data (*46*). **(I)** Analysis by ChIP-qPCR of Ser5-RNAPII binding at PRC2-target promoter regions (*Dach1, Sox7, Ascl1, Pax3, Msx1, Nkx2-2*) in G1 (black), S (grey) and G2-M (red) assayed by ChIP-qPCR. Active (*Oct4, Nanog*) and hypermethylated (*Myf5, λ-5*) gene promoters were used as controls. Mean ± SEM of three experiments are shown. **(J)** Venn diagram showing the overlap between HC-bivalent genes overtly repressed in G2-M (FC>1.5) and genes displaying accumulation of Ser5-RNAPII at their promoter region in G2-M (cluster I in Fig. 3g). **(K)** Linear regression analysis of the binding signals of Ezh2 and Ser5-RNAPII at HC-bivalent promoters (−0.5 kb to +1.5 kb relative to TSS) at indicated cell cycle phases. **(L)** Histogram displaying the percentage of genes showing increased (G1>G2-M) or decreased (G1<G2-M) binding of Ezh2 (FC>2), Jarid2 (FC>1.3), Ser5-RNAPII (FC>2), EPOP (FC>1.5), EloB (FC>1.5) and RNA production (FC>1.5) during cell cycle transition. Genes that showed no difference between analysed phases were excluded to calculate the percentage. **B, F, I.** * marks statistically significant differences.

To address whether recruitment of Jarid2-Ezh2 and firmer repression of bivalent genes during S-G2-M was associated to changes in the activity of RNA Polymerase II (RNAPII), we analysed binding of RNAPII phosphorylated in Ser5 (Ser5-RNAPII), typically associated with transcriptionally pausing and gene poising of bivalent genes in mESCs (*28*). We found augmented accumulation of Ser5-RNAPII around the TSS of HC-bivalent genes in S, G2-M as compared to G1 (Fig. 4D-F and Fig. S4E, F). Increased binding of Ser5-RNAPII was evident for most bivalent promoters (1324 out of 1677, 78.9%) genes, (see cluster I in Fig. 4G) and at individual candidate bivalent genomic domains (i.e. *Hoxd* gene cluster) (Fig. 4H). ChIP-qPCR analysis of a subset of PRC2-target genes further confirmed gradual accumulation of Ser5-RNAPII during S, G2-M compared to G1 (Fig. 4I). Most of HC-bivalent genes (592 out of 715, 82.7%) that showed reduced RNA synthesis in G2-M displayed accumulation of Ser5-RNAPII at their promoter region (Fig. 4J), suggesting that reduced production of RNA is coupled to RNAPII pausing at the promoters of bivalent genes during S-G2-M. Importantly, increased recruitment of Ezh2 (or Jarid2) during S, G2-M correlated with accumulation of Ser5-RNAPII at target promoters at all phases of the cell cycle and it became more evident during S, G2-M (Fig. 4K and Fig. S4G). Taken together (Fig. 4L), these observations indicate that increased recruitment of Jarid2-Ezh2 to target genes is associated to reduced binding of EPOP and EloB, pausing of transcription by RNAPII and reduced production of leaky RNA during S, G2-M.

### Cell-cycle-dependent regulation of PRC2 is more evident at the promoter of developmental transcription factors

We reasoned that because cell-cycle-dependent regulation of bivalent genes involved changes in recruitment of Jarid2, Ezh2, EPOP, EloB and Ser5-RNAPII, bivalent genes that are common targets of these proteins in asynchronous mESCs might display more accused regulation across cell cycle. Strikingly, we found that a set of 390 bivalent genes co-bound by these factors (common target genes) (Fig. 5A) is very significantly enriched for transcription factors and DNA binding proteins (162 out of 390 genes, p-value: 7.6 E-94), in contrast to the remaining 991 genes that are enriched for protein binding and transmembrane transporters (Fig. 5B). Common target genes included key pioneering factors involved in mesoderm, ectoderm and endoderm differentiation (Fig. S5A) suggesting that cell-cycle-dependent regulation of Polycomb recruitment modulates differentiation to the three germ layers. Importantly, comparative analysis of PRC2 binding revealed that common targets are more profoundly bound by PRC2 subunits (Ezh2, Jarid2 and EPOP) and Ser5-RNAPII than remaining genes, and that their differential recruitment of PRC2 subunits, Ser5-RNAPII and RNA production across cell cycle is more accused (Fig. 5C and Fig. S5B-D). In agreement, analysis using published data of PRC2 binding in asynchronous populations of mESCs showed that recruitment of Ezh2, Jarid2, EPOP and EloB as well as Eed, Suz12, Mtf2 and H3K27me3 was higher at the promoters of common targets than at the promoters of remaining genes (Fig. S5E). In contrast, common targets were not enriched for trithorax protein MLL2 and H3K4me3 (Fig. S5E). Taken together these results indicate that the promoter regions of transcription factors that regulate cell differentiation recruit higher levels of PRC2 subunits that are prominently regulated across cell cycle.

**Figure 5.**
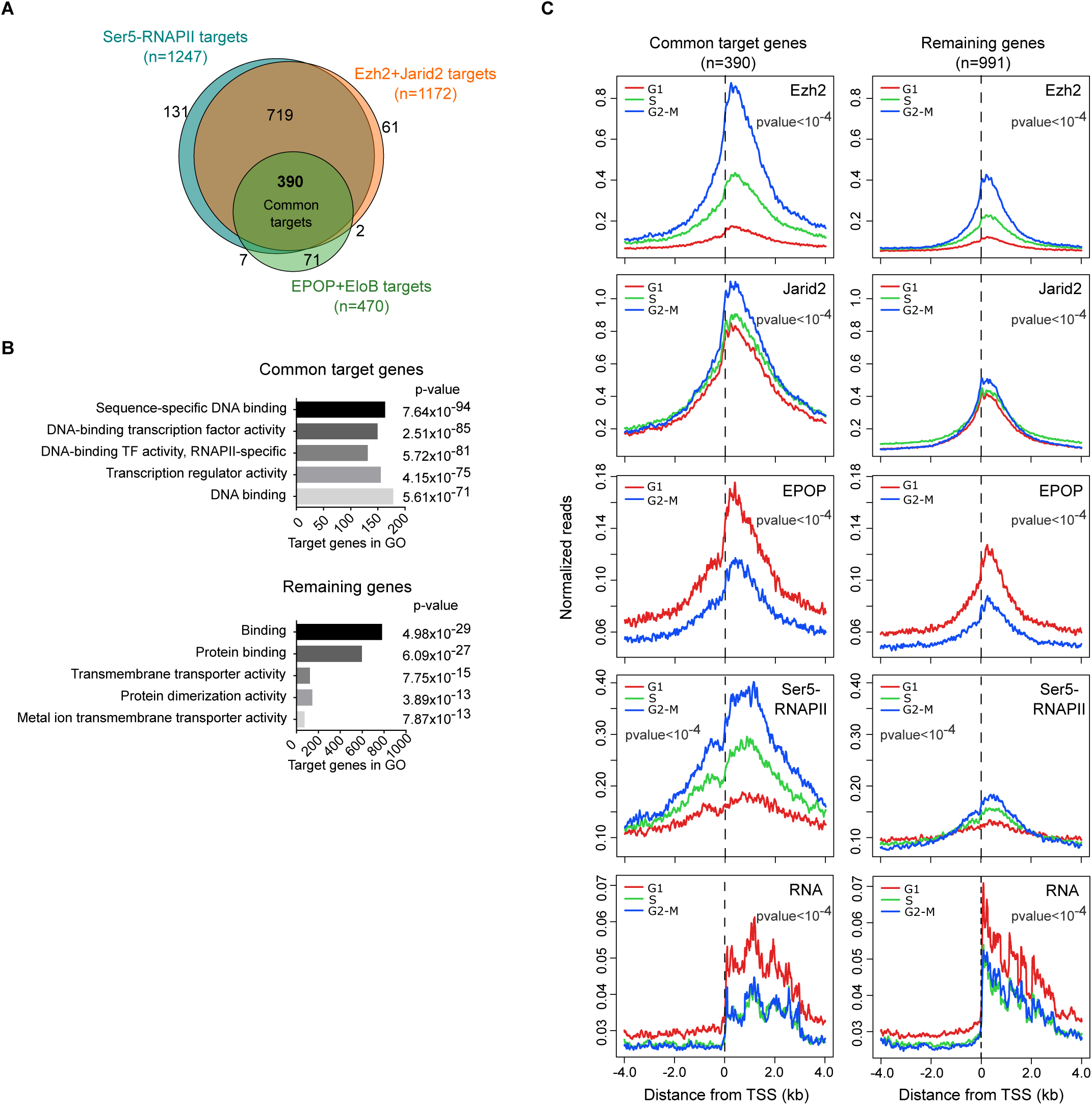
Cell-cycle-dependent regulation of PRC2 is more accused at the promoter of developmental transcription factors. **(A)** Venn diagram showing overlap between HC-bivalent genes that are targets of Ser5-RNAPII, Ezh2, Jarid2, EPOP and EloB in asynchronous populations using published (*23, 36*) and our (Jarid2) datasets. **(B)** Gene Ontology analysis of common target and remaining genes. Bars represent the number of genes that fall into indicated GO categories. p-value is shown next to each category. **(C)** Average binding profile of Ezh2, Jarid2, EPOP, Ser5-RNAPII and RNA synthesis around the TSS of HC-bivalent genes in G1 (red), S (green) and G2-M (blue) comparing common target and remaining HC-bivalent genes (as defined in Fig. 5A).

### Cell-cycle-dependent regulation of PRC2 is hindered in Jarid2 -/- mESCs

We next tested how the lack of the PRC2-recruiter Jarid2 affected cell-cycle-dependent regulation of PRC2. We derived Fucci-Jarid2 -/- by introducing the Fucci reporter system into previously derived Jarid2 -/- mESCs (*20*) (Fig. 6A). Jarid2 depleted mESCs display reduced binding of core PRC2 subunits and Ser5-RNAPII (*18, 20*) to bivalent promoters, but they show unchanged levels of EPOP binding at target genes (*23*). In fitting, analysis of RNA expression and recruitment of Ser5-RNAPII in Fucci-Jarid2 fl/fl compared to Fucci-Jarid2 –/- mESCs, revealed that the lack of Jarid2 results in evident de-repression of RNA synthesis coupled with loss of paused Ser5-RNAPII from gene promoters in G2-M but only subtle genes in G1 (Fig. 6B, C). Analysis by ChIP-qPCR further confirmed that accumulation of paused Ser5-RNAPII at bivalent genes in G2-M is visibly reduced in Jarid2 -/- compared to parental mESCs (Fig. 6D). Thus, these results demonstrate that increased recruitment of Jarid2 and Ezh2 to bivalent genes during S, G2-M results in pausing of RNAPII and reduced production of RNA.

**Figure 6.**
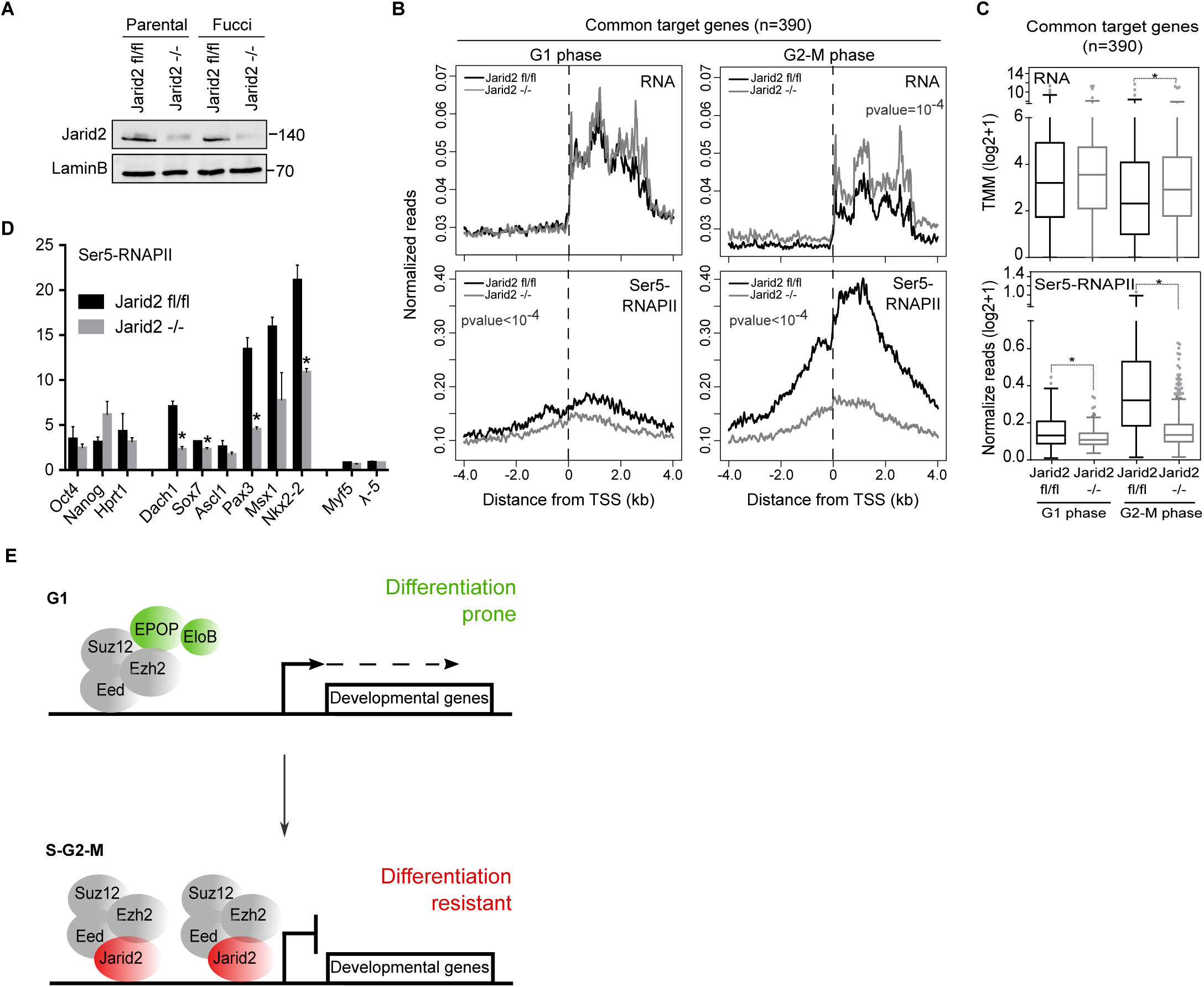
Jarid2 -/- mESCs display altered cell cycle-dependent regulation of bivalent genes. **(A)** Whole cell extracts western blots of Jarid2 in Fucci-Jarid2 fl/fl and Fucci-Jarid2 -/- mESCs. **(B)** Average RNA synthesis (top panels) and binding of Ser5-RNAPII (bottom panels) around the TSS of common target bivalent genes in G1 (left panels) and G2-M (right panels) in *Jarid2* fl/fl (black lines) and *Jarid2* -/- (grey lines) mESCs. **(C)** Quantification of RNA synthesis (left) and Ser5-RNAPII binding (right) in *Jarid2* fl/fl and *Jarid2* -/- cells in G1 and G2-M at common target genes. **(D)** Analysis by ChIP-qPCR comparing the binding of Ser5-RNAPII to PRC2-target promoter regions (*Dach1, Sox7, Ascl1, Pax3, Msx1, Nkx2-2*) in *Jarid2* fl/fl (black bars) and *Jarid2* -/- (grey bars) in G2-M phase. Active (*Oct4, Nanog*) and hypermethylated (*Myf5, λ-5*) gene promoters were used as controls. Mean ± SEM of three experiments are shown. * marks statistically significant differences. **(E)** Schematic diagram of observations described in this manuscript. **C, D.** * marks statistically significant differences.

## Discussion

Our results show that recruitment of PRC2 complexes to target promoters is regulated across cell-cycle and indicate that PRC2.1 complexes are preferentially recruited in G1, while binding of PRC2.2 is favoured in S, G2-M (Fig. 6E). This is supported by our ChIP-seq analysis revealing augmented binding of EPOP and EloB (part of PRC2.1) to target promoters in G1 as opposed to increased binding of Jarid2 (part of PRC2.2) during S, G-M phases. Importantly, we found that binding of the catalytic subunit Ezh2 is dramatically increased in S, G2-M compared to G1, indicating that cell-cycle regulation is not exclusive of accessory subunits but is also happening for the core catalytic subunit of the complex. This is in fitting with previous reports suggesting a cell-cycle-dependent regulation of Ezh2 binding by CDK1 and CDK2 phosphorylation (*29*). Jarid2 activity is known to favour high levels of PRC2 binding (*18-22*) while EPOP activity promotes the opposite (*23*). Thus, reduced recruitment of Ezh2 in G1 is probably a consequence of the accumulation of EPOP-PRC2 (PRC2.1) at the expense of Jarid2-PRC2 (PRC2.2) complexes in this cell cycle phase. Importantly, we showed that nascent RNA produced in G1 is still in the range of leaky transcription rather than full activation of target genes, suggesting that although the amount of Ezh2 bound to promoters of bivalent genes in G1 is lower than in S, G2-M, it is probably enough to maintain significant gene repression of target genes. Notwithstanding, repression of Ezh2-target genes in G1 might also be dependent on Ezh2-independent mechanisms (*11*) including the activity of Ezh1 or PRC1 complexes.

We found that recruitment of Jarid2-PRC2 (PRC2.2) during S, G2-M phases leads to more robust gene repression of target bivalent genes; reduced production of RNA coupled with increased binding of paused Ser5P-RNAPII in S, G2-M that is lost upon Jarid2 depletion. These observations, together with the known role of EPOP in recruiting EloB and promoting leaky transcription at bivalent genes (*23, 30*) support that recruitment of EPOP-PRC2 (PRC2.1) to bivalent promoters in G1 favours a transcriptional permissive chromatin setup. Importantly, accumulation of Ser5P-RNAPII is temporally coincident with recruitment of PRC2.2 during S, G2-M. Because simultaneous binding of PRCs and Ser5-RNAPII at bivalent gene promoters is a key defining feature of transcriptional priming, our results highlight a previously unanticipated regulation of gene priming within the pluripotent cell cycle. In particular, our results demonstrate that gene priming is evident during S, G2-M (high levels of Ezh2 and Ser5-RNAPII) but it is dismantled during G1 (low level of Ezh2 and Ser5-RNAPII), rather than being maintained until the exit of the pluripotent cell cycle.

Asynchronous populations of pluripotent cells display heterogenous expression of genes and cell differentiation ability (*2-4*). This is partly because G1-cells display elevated expression of developmental regulators (*8*) and they are more prone to activate lineage specific genes in response to differentiation cues (*5, 6*). We found that mESCs in G1 display a pro-activation chromatin setup characterized by enhanced binding of the PRC2 activating subunits EPOP and EloB, together with reduced binding of Jarid2 and Ezh2. We showed that this leads to alleviation of transcriptional repression in G1. In fitting, it has been reported that hESCs display increased H3K4me3 at the promoter of bivalent genes during G1 as compared to S, G2-M (*31*). Thus, given that de-repression of PRC2-target genes is a critical early event during cell differentiation (*32*), transient alleviation of Polycomb repression in G1 is a key observation to explain why pluripotent cells in G1 are more sensitive to differentiation signals.

Overall, in this study we showed for the first time that Polycomb complexes subunit configuration is controlled by cell-cycle dependent mechanisms in mESCs, and that this type of regulation is functionally relevant in the context of pluripotent cell differentiation. Because the general principles underlying Polycomb function are widely conserved from flies to humans (*11*), our discovery will probably be relevant for other model systems in which the regulation of gene expression by Polycomb needs to be coordinated with DNA replication and cell division, including adult stem cells and tumour cells (*33*). Additionally, PRCs can also regulate gene activity in somatic G1-arrested cells (*33*) and thus, in the future, it will be interesting to see to what extend differences in the regulation by Polycomb proteins in different model systems are a consequence of their different cell-cycle configuration.

## Methods

### ESCs growth, Fucci and flow cytometry

Stable Fucci mES cell lines (background 129/Sv/C57BL/C6) were generated for parental and *Jarid2* knockout mESCs (*20*) by transfecting the ES-FUCCI plasmid (*34*) (Addgene repository #62451). mESCs expressing mCherry:hCdt and Citrine:Geminin were cultured in 5% CO2 at 37 °C on 0.1% gelatin-coated dishes in DMEM KO (Gibco) media supplemented with 10% FCS, leukaemia-inhibiting factor (LIF), penicillin/streptomycin (Gibco), L-glutamine (Gibco), 2-mercaptoethanol (Gibco) and hygromycin B (InvivoGen) as described previously (*35*). Upon trypsinization and resuspension in sorting buffer (PBS, 2% FCS, EDTA 2mM and LIF) at 4°C, Fucci-mESCs were cell sorted in an Aria Fusion flow cytometer equipped with 488 and 561 lasers to discriminate cells expressing citrine (516 nm/529 nm) and mCherry (587nm/610 nm). Sorted cells were counted and 1 million cells were used typically for downstream genome-wide analysis. Cell cycle profile of sorted cell populations was routinely checked by propidium iodide staining followed by flow cytometry.

### Gene promoter classification

The list of bivalent genes (promoter positive for H3K4me3 and H3K27me3) (3753 genes) was described previously (*36*). HC-Bivalent gene list (1678 genes) was obtained by extracting bivalent genes consistently described in three different studies (*36-38*). Active genes (1557 genes) were determined using published data (*36*): genes positive for H3K4me3, bound by phosphorylated (Ser5, Ser2, Ser7), hypophosphorylated RNAPII (8WG16 antibody), with RNA expression higher than 20 FPKM and negative for PRC2 binding (Ezh2, Suz12 and H3K27me3). Hypermethylated promoters in mESCs (more than 80% CpG methylation) were identified using published bisulphite sequencing data (*39*) and crossed analysed with published data (*36*) to identify hypermethylated promoters transcriptionally silent (less than 1 FPKM) and not bound by PRC2 (Ezh2, Suz12 and H3K27me3) (656 genes). See supplementary data for complete gene lists.

### Chromatin immunoprecipitation qPCR and sequencing

ChIP assays for Ser5-RNAPII, Ezh2, Jarid2 and EPOP were performed as described previously (*36*) with minor modifications: Typically, 1 million cell-cycle-sorted cells were resuspended in 37°C complete media (200 ul per million cells) and incubated in a rotating platform for 12 min with 1% formaldehyde at room temperature. To stop the reaction, glycine was added to a final concentration of 125 mM. Swelling and sonication buffers (protease and phosphatase inhibitors supplemented) were used at 4°C in a proportion of 0.5 ml per million cells. Ezh2, Jarid2 and EPOP ChIPs were carried without using bridge antibody. Chromatin and antibody were incubated in a rotating wheel at 4°C overnight. Protein G magnetic beads (Dynabeads, Invitrogen) were then added and incubated for 5 hours. Washes were carried out for 5 min at 4°C with 1 ml of the following buffers: 1 x Sonication Buffer, 1 x Wash Buffer A (50 mM HEPES pH 7.9, 500 mM NaCl, 1 mM EDTA, 1% Triton X-100, 0.1% Na-deoxycholate, 0.1% SDS), 1 x Wash Buffer B (20 mM Tris pH 8.0, 1 mM EDTA, 250 mM LiCl, 0.5% NP-40, 0.5% Na-deoxycholate), 2 x TE Buffer pH 8. ChIP of EloB was performed as above but adding a chromatin double-crosslink step as describe in (*23*) with minor modifications: Cells were resuspended in PBS at 4°C after flow cytometry sorting and incubated with ChIP Crosslink Gold (Diagenode #C01019027, 0.8 ul in 200 ul of PBS per 1 million cells) in a rotating platform for 30 min at room temperature. After one wash step chromatin was crosslinked with 1% formaldehyde for 10 min at room temperature. ChIP-qPCR of Ezh2, EPOP and Ser5-RNAPII and quality control of all immunoprecipitated DNA samples were tested by qPCR using GoTaq® qPCR Master Mix (Promega) with a QuantStudio 6 Flex Real-time PCR System (Applied Biosystems). Enrichment was calculated relative to 1% input for all ChIPs except for Ser5-RNAPII that was normalized by loading the same amount of DNA as described previously (*36*). Details of antibodies and primers used are available in Supplementary methods.

Libraries of immunoprecipitated DNA in Ezh2, Jarid2, EPOP and Elob ChIP-seqs were generated from 1-5 ng of starting DNA with the NEBNext Ultra DNA Library Prep kit for Illumina (#7370) according to manufacturer’s instructions at CRG Genomics Core Facility (Barcelona) and sequenced using a HiSeq 2500 Illumina technology. Library of Ser5-RNAPII ChIP-seq was performed using the NextFlex ChIP-seq kit (Bioo #NOVA-5143-01) starting with 4 ng of immunoprecipitated DNA and sequenced at Centre for Genomics and Oncological Research (Genyo, Granada) using Illumina technology (NextSeq 500) according to the manufacter’s instructions. 20-30 million reads (50 bp single-reads) were obtained for each library.

Reads were aligned and quantified using STAR 2.5.2 (*40*) against GENCODE NCBI m37 (mm9) genome. Samtools 1.3.1 (*41*) was used to discard alignments with a quality score < 200 in order to remove multi-mapping reads. Finally, we used BamCompare from deepTools suite (*42*) to create bigwig files with the signal normalized by Reads-per-million (RPM) and against an input sample. Peak calling was performed with MACS2. Data mining of publicly available ChIP-seq datasets (Eed, Suz12, Ezh2, H3K27me3, Jarid2, EPOP, EloB, Mtf2, Mll2, H3K4me3) were treated the same way. CoverageView (Coverage visualization package for R. R package version 1.20.0.) was used to calculate coverage around Transcription Start Sites (TSS).

Average normalized reads (RPM) for a genomic window of −0.5 kb to +1.5 kb relative to TSS for each analysed promoter was calculated and represented as boxplots and were subjected to clustering using Cluster 3.0 followed by the Java TreeViewer software. Log2 of binding values relative to the average were used. Average binding plots were generated by counting normalize reads every 10 bp. In heatmaps analyses of reads density in ChIP-seq experiments we log2-transformed RPMs and trimmed these values between the minimum 5th percentile and the maximum 95th percentile. To compare different samples, genes were ranked according to G2-M (Ezh2, Jarid2 and Ser5-RNAPII) or G1 (EPOP and EloB). Gene Ontology analysis was performed using the Gene Ontologie knowledge database (geneontology.org).

### 4-thiouridine-tagging sequencing (4sU-seq)

4sU-seq experiments were carried out as described in (*27*) with minor modifications: cells were treated with 4-thiouridine (500 uM) during 1-hour 37°C before trypsinization and flow cytometry sorting. 1 million sorted-cells were resuspended in 500 ul of Trizol (Invitrogen). Isolated RNA was Bioanalyzed (Agilent) and was subjected to qPCR analysis as quality control. 4sU-RNA and total RNA samples were retrotranscribed using RevertAid RT Reverse Transcription kit (Thermo Fisher #K1691), treated with Dnase I (Thermo Fisher #18068015) and were subjected to qPCR using primers designed to specifically amplify unspliced RNA (contiguous exon-intron sequences) and total RNA (exon sequences). Strand specific 4sU-seq libraries were generated using 75 ng of 4sU-RNA and the NextFlex Rapid Directional RNA-seq kit (Bioo Scientific #NOVA-5138-07) according to manufacturer’s instructions. Libraries were quantified by NanoDrop 2000 (Thermo Fisher) and Bioanalyzed. 50 million 75 bp paired-end reads per sample were sequenced using Illumina technology (NextSeq 500) at Centre for Genomics and Oncological Research (Genyo).

After quality control, we used SortMeRNA 2.1 software (*43*) to filter out rRNA reads. We aligned and quantified filtered reads with STAR 2.5.2 (*40*) using GENCODE NCBI m37 (mm9) as the reference genome. We used Samtools 1.3.1 (*41*) to remove alignments with a quality score < 200 in order to discard multi-mapping reads. Finally, we used BamCompare from deepTools suite (*42*) to create bigwig files with normalized signal in Reads-per-million (RPM) and with positive values for forward strand and negative values in reverse strand. In order to normalize gene expression values, we applied trimmed mean of M values (TMM) (*44*) method with NOISeq package (*45*).

### Western Blot analysis

Whole extracts were prepared for 1 million cells after flow cytometry sorting. Cells were pelleted and resuspended in 50 ul of PBS and 50 ul of 2X Laemly Buffer (0.1 M Tris pH 6.8, 2% SDS, 5% glycerol) supplemented with protease and phosphatase inhibitors (1X EDTA-Free inhibitor cocktail (Roche), 1 mM PMSF, 5 mM NaF, 2 mM Na3VO4). Western was carried out using standard procedures.

### Statistical analysis

Statistical analyses were carried out using R 3.5.1. In boxplots, whiskers denote the interval within 1.5x the interquartile range and p-values were calculated using Mann-Whitney test (significant differences p-value<0.0001). Average mapped reads around the TSS was carried out using ANOVA, comparing all samples in a window of −0.5 to +1.5 kb from TSS (significant differences p-value<0.0001). ChIP-qPCR statistical analysis was carried out for triplicates and using T-Student test (significant differences p<0.05).

### Data access

Datasets are available at GEO-NCBI with accession number GSE128851 (https://www.ncbi.nlm.nih.gov/geo/query/acc.cgi?acc=GSE128851) with a temporal private token “grsjscwuvruzdmb” that will be released upon publication acceptance.

## Supporting information

ChIP-seq results

Gene lists used in analyses

## Disclosure declaration

Authors declare no competing interests.

## Author contributions

DL designed the study. HGA and DL wrote the manuscript. HGA, AG designed, performed and analysed experiments. JM and PC carried out bioinformatic analysis.

## Acknowledgments

We are very grateful to Ana Pombo, Emily Brookes and Lars Dölken for sharing in house protocols and technical advice for Ser5-RNAPII and 4sU-seq analyses. To Luciano Di Croce for sharing EPOP and EloB home-made antibodies and providing scientific advice. To core facilities in GENYO and in particular to the flow cytometry and genomics units. We also thank the genomics unit at the CRG for assistance with ChIP-seq experiments. This study was supported by the spanish ministry of economy and competitiveness (SAF2013-40891-R; BFU2016-75233-P) and the andalusian regional government (PC-0246-2017). David Landeira is a Ramón y Cajal researcher of the Spanish ministry of economy and competitiveness (RYC-2012-10019).

## Figure Legends

**Figure S1.**
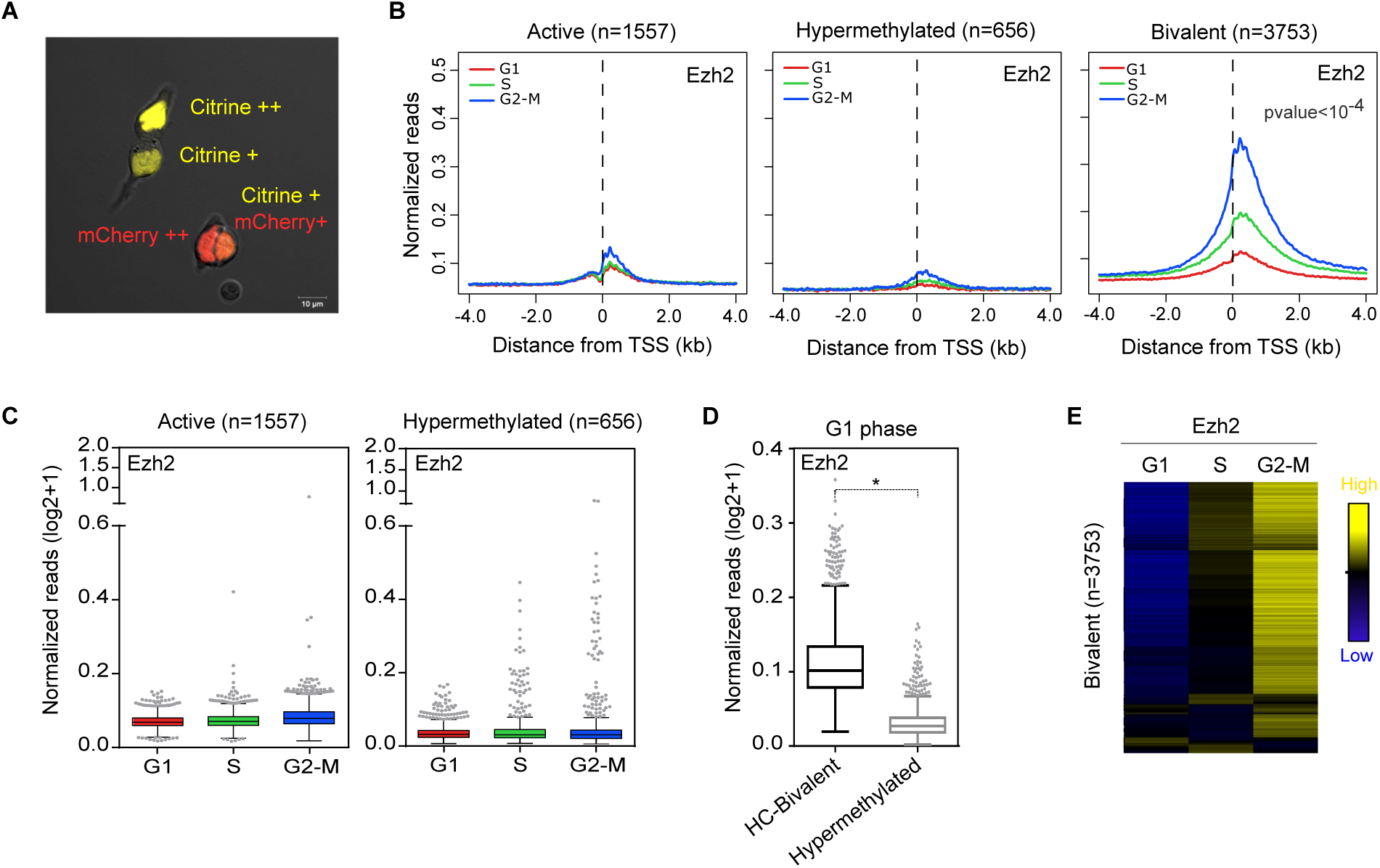
Differential recruitment of the PRC2 catalytic subunit Ezh2 across cell cycle. **(A)** Fluorescence microscopy image showing Fucci-mESCs expressing expected combinations and intensities of Citrine:Geminin and mCherry-hCdt1. **(B)** Average binding profile of Ezh2 around the TSS of active, hypermethylated and bivalent (as defined in (*36*)) genes in G1 (red), S (green) and G2-M (blue). **(C)** Quantification of Ezh2-binding signal at the promoter regions (−0.5 kb to +1.5 kb relative to TSS) of active and hypermethylated genes in indicated cell cycle phases. **(D)** Boxplot quantification of Ezh2-binding signal at the promoter regions of HC-bivalent (black) and hypermethylated (grey) promoters in G1. * marks statistically significant differences. **(E)** Hierarchical clustering analysis of binding of Ezh2 to the promoter region (−0.5 kb to +1.5 kb relative to TSS) of bivalent genes (as defined in (*36*)) at indicated phases of the cell cycle. Binding relative to the average is presented.

**Figure S2.**
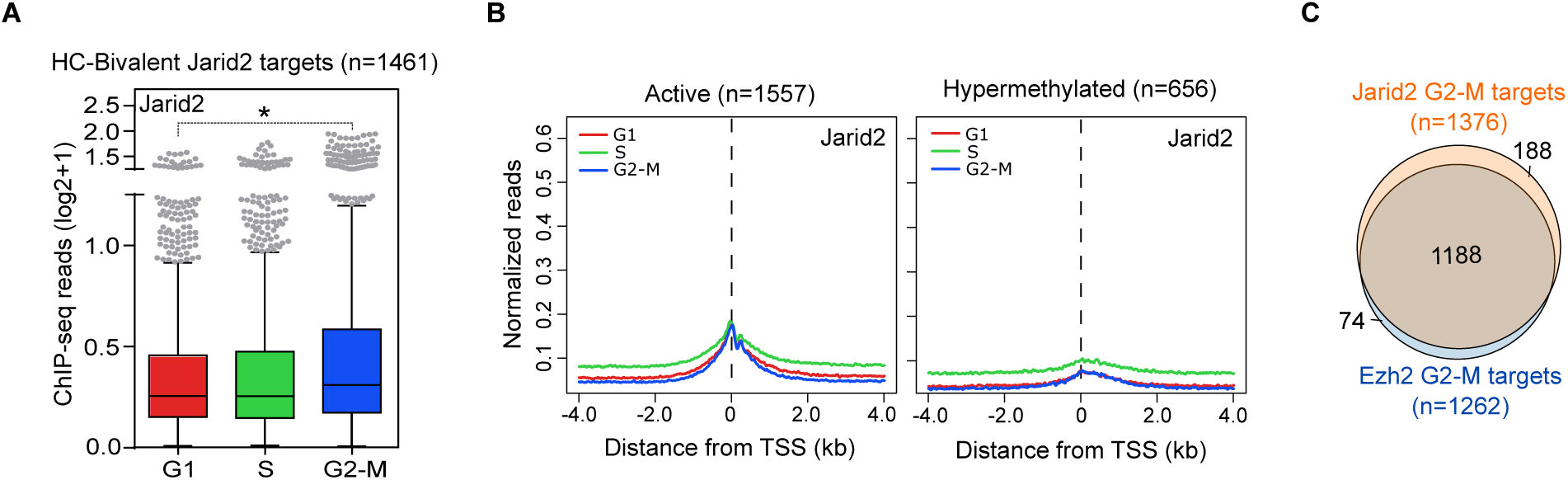
Cell-cycle-dependent regulation of Jarid2. **(A)** Boxplot of Jarid2-binding signal at the promoter regions (−0.5 kb to +1.5 kb relative to TSS) of HC-bivalent genes in indicated cell cycle phases. * marks statistically significant differences. **(B)** Average binding profile of Jarid2 around the TSS of active and hypermethylated gene promoters in G1 (red), S (green) and G2-M (blue). **(C)** Venn diagram showing overlapping between Jarid2 and Ezh2 target genes in G2-M.

**Figure S3.**
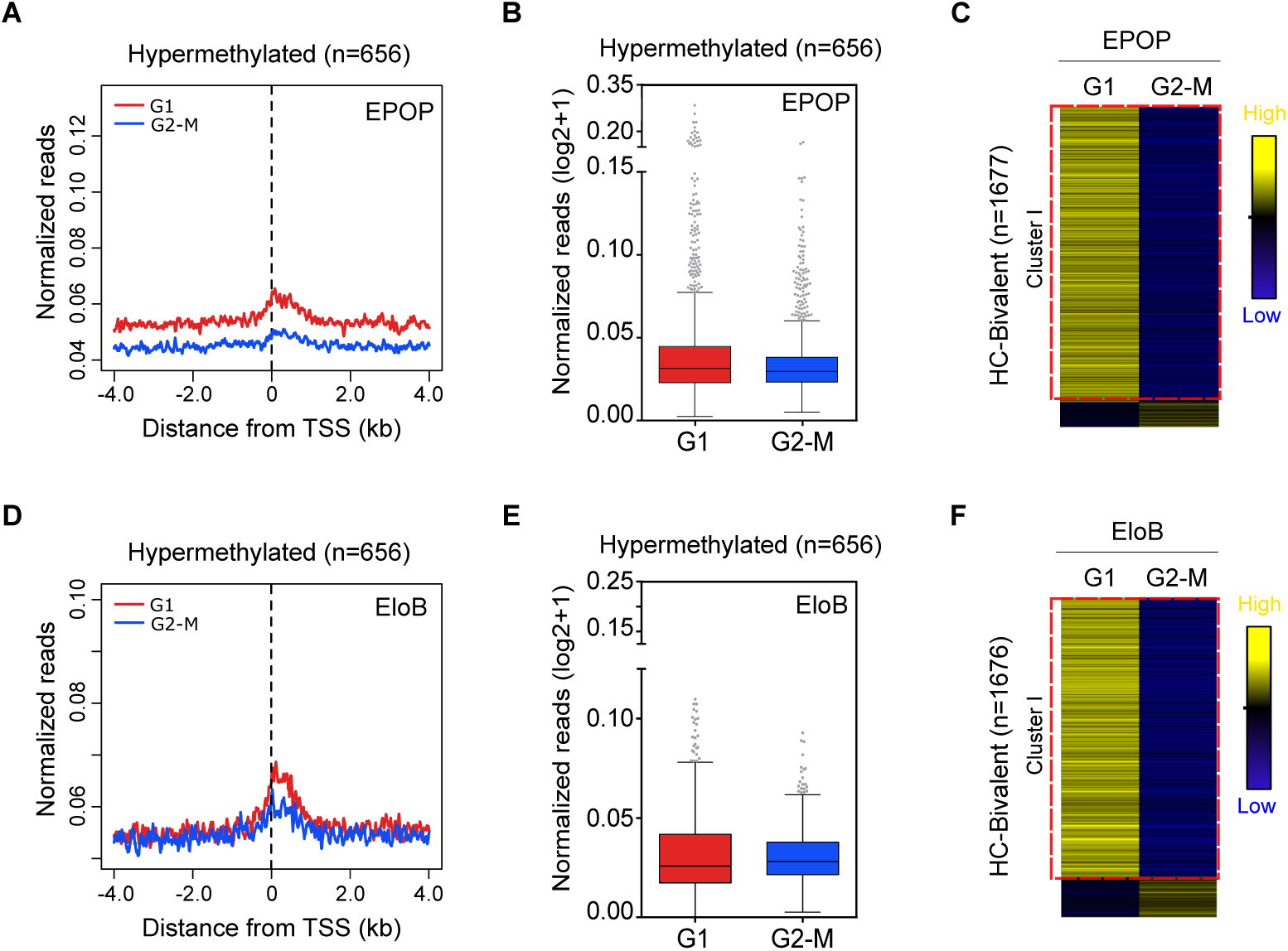
Binding of EPOP and EloB to bivalent promoters is regulated across cell cycle. **(A)** Average binding profile of EPOP around the TSS of hypermethylated gene promoters in G1 (red) and G2-M (blue). **(B)** Boxplot of EPOP-binding signal at the promoter regions (−0.5 kb to +1.5 kb relative to TSS) of hypermethylated genes in indicated cell cycle phases. **(C)** Hierarchical clustering analysis of binding of EPOP to the promoter region (−0.5 kb to +1.5 kb relative to TSS) of HC-bivalent genes at indicated phases of the cell cycle. Binding relative to the average value is presented. **(D)** Average binding profile of EloB around the TSS of hypermethylated gene promoters in G1 (red) and G2-M (blue). **(E)** Boxplot of EloB -binding signal at the promoter regions (−0.5 kb to +1.5 kb relative to TSS) of hypermethylated genes in indicated cell cycle phases. **(F)** Hierarchical clustering analysis of binding of EloB to the promoter region (−0.5 kb to +1.5 kb relative to TSS) of HC-bivalent genes at indicated phases of the cell cycle.

**Figure S4.**
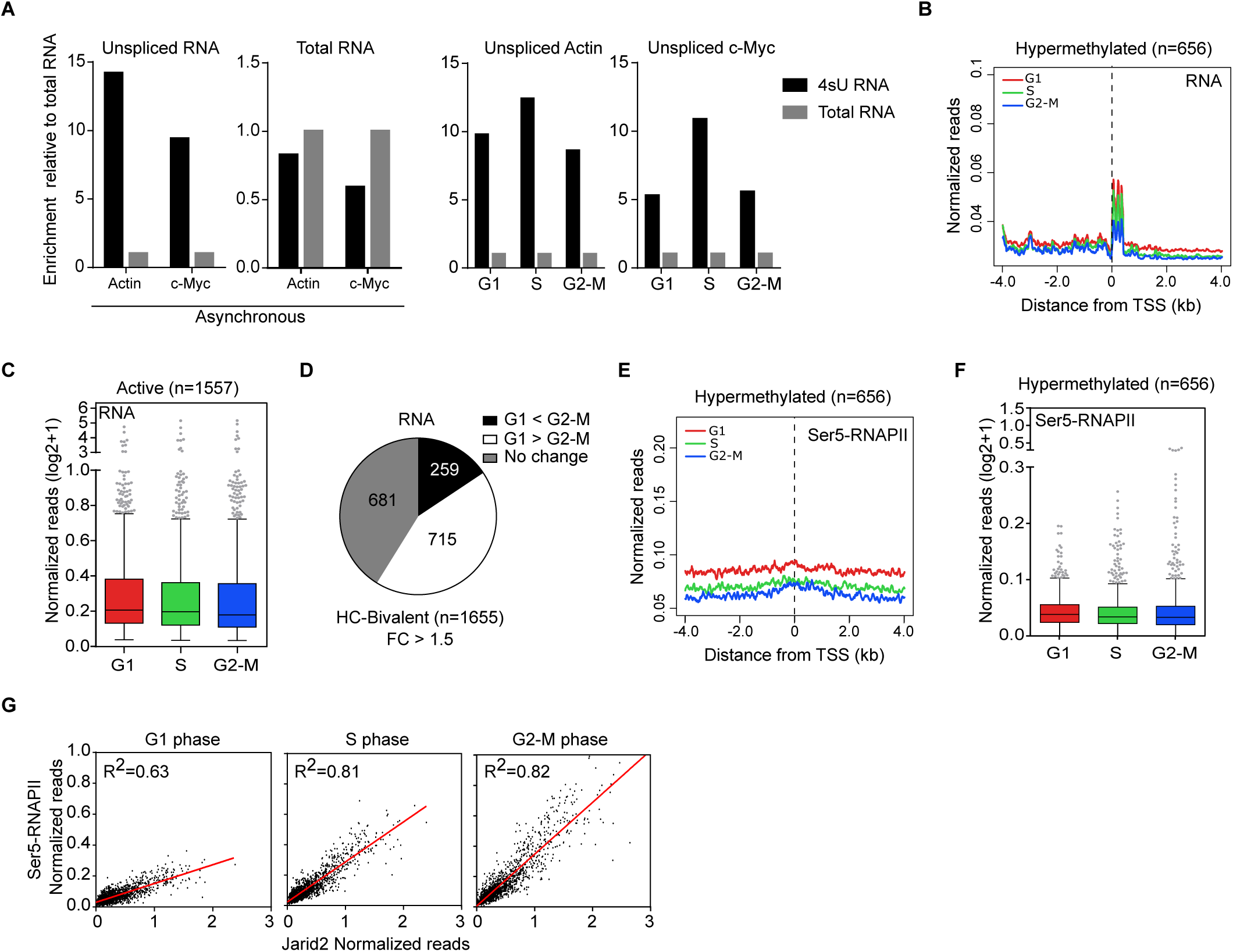
Transcriptional activity at PRC2-target genes is attenuated during S, G2-M. **(A)** Histogram comparing the enrichment of unspliced RNA (Actin and c-Myc) in 4sU RNA fraction in asynchronous (left panel) and cell-cycle sorted (right panel) cells by RT-qPCR. **(B)** Average RNA production from hypermethylated promoters in G1 (red), S (green) and G2-M (blue). **(C)** Quantification of Ser5-RNAPII-binding signal at the promoter regions (−0.5 kb to +1.5 kb relative to TSS) of active genes in indicated cell cycle phases. **(D)** Diagram representing the number of HC-bivalent genes showing changes in RNA expression between G1 and G2-M (FC > 1.5). **(E)** Average binding profile of Ser5-RNAPII around the TSS of hypermethylated gene promoters in G1 (red), S (green) and G2-M (blue). **(F)** Quantification of Ser5-RNAPII-binding signal at the promoter regions (−0.5 kb to +1.5 kb relative to TSS) of hypermethylated genes in indicated cell cycle phases. **(G)** Linear regression analysis of the binding signals of Jarid2 and Ser5-RNAPII at HC-bivalent promoters (−0.5 kb to +1.5 kb relative to TSS) at indicated cell cycle phases.

**Figure S5.**
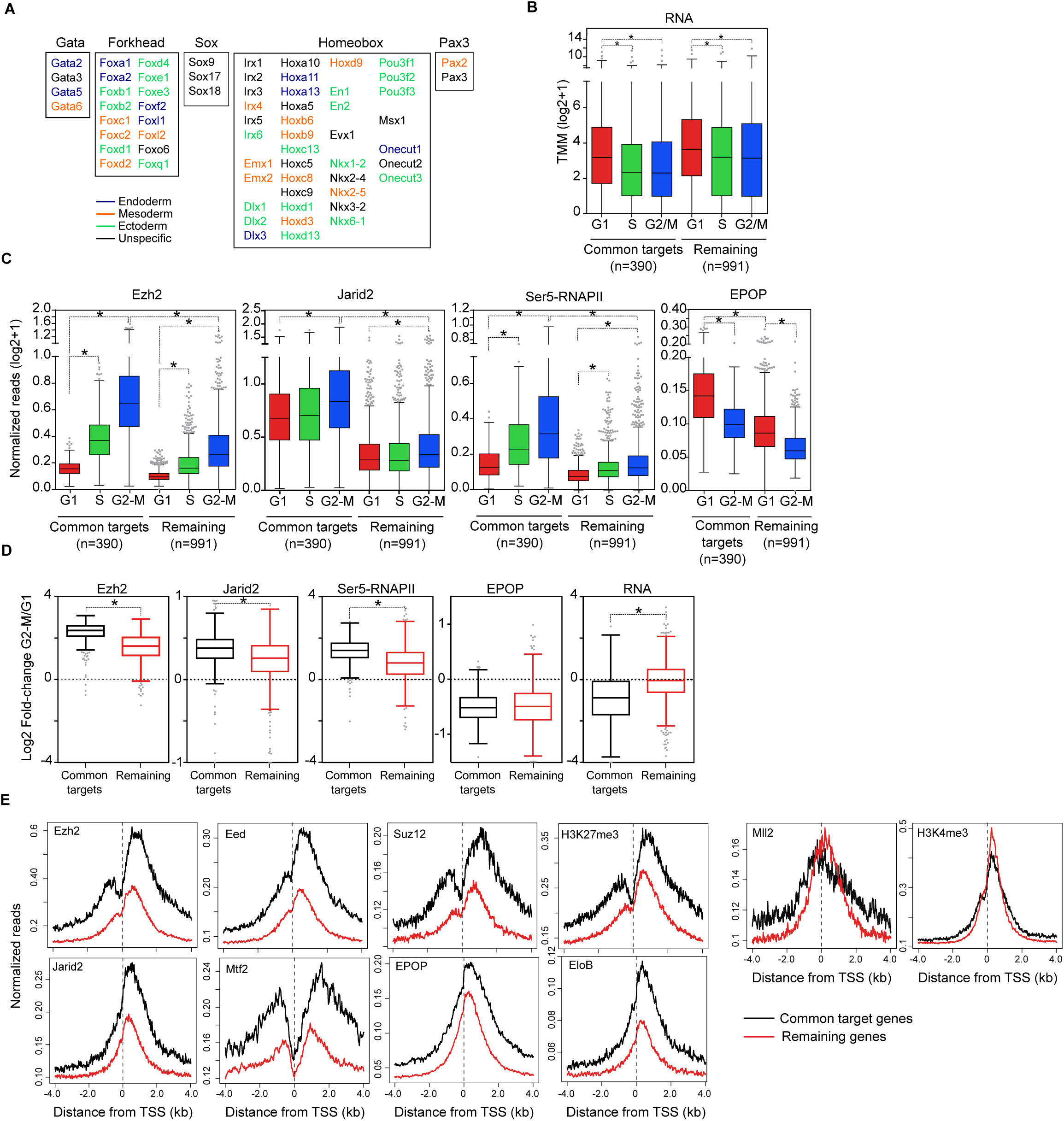
PRC2-target promoters that are regulated across cell cycle display higher levels of PRC2 binding. **(A)** List of transcription factors identified in the subset of common targets. Reported association to endoderm (blue), mesoderm (orange) and ectoderm (green) is indicated. **(B)** Quantification of RNA expression (TMM normalized) of common targets and remaining bivalent genes in indicated cell cycle phases. **(C)** Quantification of the binding of Ezh2, Jarid2, Ser5-RNAPII and EPOP at the promoter regions (−0.5 kb to +1.5 kb relative to TSS) of common targets and remaining bivalent genes in indicated cell cycle phases. **(D)** Boxplots showing the fold change of RNA synthesis (TMM) and binding of Ezh2, Jarid2, Ser5-RNAPII at promoter regions (−0.5 kb to +1.5 kb relative to TSS) between G2-M and G1 in common targets (black boxplots) and remaining (red boxplots) bivalent genes. **(E)** Plots comparing the average binding profiles of Ezh2, Eed, Suz12, H3K27me3, Jarid2, Mtf2, EPOP, EloB, MLL2 and H3K4me3 in asynchronous populations at common target (black lines) and remaining (red lines) bivalent genes.

